# Cognate epitope recognition by bovine CD4 and CD8 T cells is essential for primary expansion of antigen-specific cytotoxic T-cells following ex vivo stimulation with a candidate *Mycobacterium avium subsp. paratuberculosis* peptide vaccine

**DOI:** 10.1101/555672

**Authors:** Gaber S. Abdellrazeq, Lindsay M. Fry, Mahmoud M. Elnaggar, John P. Bannantine, David A. Schneider, William M. Chamberlin, Asmaa H. A. Mahmoud, Kun-Taek Park, Victoria Hulubei, William C. Davis

## Abstract

Studies in cattle show CD8 cytotoxic T cells (CTL), with the ability to kill intracellular bacteria, develop following stimulation of monocyte-depleted peripheral blood mononuclear cells (mdPBMC) with conventional dendritic cells (cDC) and monocyte-derived DC (MoDC) pulsed with MMP, a membrane protein from *Mycobacterium avium* subsp. *paratuberculosis* (*Map*) encoded by *MAP2121c*. CTL activity was diminished if CD4 T cells were depleted from mdPBMC before antigen (Ag) presentation by cDC and MoDC, suggesting simultaneous cognate recognition of MMP epitopes presented by MHC I and MHC II molecules might be essential for development of CTL activity. To clarify whether cognate recognition is essential for CTL development, studies were conducted with mdPBMC cultures in the presence of monoclonal antibodies (mAbs) specific for MHC class I and MHC class II molecules. The CTL response of mdPBMC to MMP-pulsed DC was completely blocked in the presence of mAbs to both MHC I and II molecules and also blocked in the presence of mAbs to either MHC I or MHC II. The results demonstrate CD4 T-cell help is essential for development of a primary CTL response to MMP, and indicate that cognate recognition is required for delivery of CD4 T-cell help during priming. Of importance, the findings provide support for the importance of CD4 and CD8 T-cell cognate antigen recognition in eliciting CTL responses to vaccines against intracellular pathogens. The methods described herein can be used to elucidate the intracellular interactions between lymphocytes and DC in humans and cattle.

## Introduction

*Mycobacterium avium* subspecies *paratuberculosis* (*Map*) is a higher-order bacterial pathogen with a broad host range that includes livestock and humans (1, 2). Similar to *M. tuberculosis* and *M. bovis*, initial infection leads to development of a persistent infection under immune control. In some cases, immune control may become compromised, leading to granulomatous ileitis, malabsorption, wasting, and death (3).

Paratuberculosis (Ptb), also referred to as Johne’s disease (JD) in ruminants, is a significant cause of livestock morbidity and mortality worldwide. The increased incidence of *Map* infection in ruminants has been accompanied by an increased prevalence of *Map* infection in humans. Some infected individuals have developed granulomatous ileitis similar to that observed in JD-infected ruminants. Interestingly, the lesions and the resultant intestinal illness are often observed in patients with Crohn’s disease (CD) (2, 4, 5), and *Map* has been cultured from numerous patients with CD. Such observations have increased interest in developing methods to limit *Map* infection in livestock, thereby reducing the risk for human exposure.

With this objective in mind, we developed a candidate peptide-based vaccine for *Map* that elicits development of CD8 cytotoxic T cells capable of killing intracellular bacteria (6). Following inoculation with wild type (WT) *Map* or *Map* deletion mutants (*Map*/Δ*relA* or *Map*/Δ*pknG* (7, 8)), calves develop both CD4 and CD8 T-cell proliferative responses to *Map* soluble antigens and Johnin (purified protein derivative, PPD made from *Map*), both before and after subsequent challenge with *Map* (9). Interestingly, only the *Map*/Δ*relA* mutant was unable to establish a persistent infection. In addition, calves inoculated with *Map*/Δ*relA* exhibited reduced colonization by WT *Map* on subsequent challenge. These data indicated a difference in the immune response of calves to *Map*/Δ*relA* as compared to that elicited by WT *Map* (9).

The immune response of a steer inoculated with *Map/*Δ*relA* was evaluated in an attempt to understand why this mutant could not establish a persistent infection. PBMC, conventional dendritic cells (cDC) present within monocyte-depleted PBMC (mdPBMC), and monocyte-derived DC (MoDC) from a steer were pulsed with live *Map*/Δ*relA* to examine the recall response (10). Initial studies showed stimulation of the calf’s mdPBMC, with cDC and MoDC pulsed with *Map/*Δ*relA* elicited a proliferative CD4 and CD8 T cell recall response. Analysis of effector activity revealed that the responding CD8 T cells were cytotoxic, killing intracellular bacteria (6). Minimal induction of cytotoxic activity was detected in the responding CD4 T cell population. Further analysis of the recall immune response to *Map/*Δ*relA* revealed the target of the response was a 35 kDa membrane peptide known as MMP, encoded by *MAP2121c* (10). An identical CTL recall response was elicited when cDC and MoDC were pulsed with MMP (6). The recall responses to *Map/*Δ*relA* and MMP were blocked in the presence of monoclonal antibodies (mAbs) specific for MHC class I and II molecules, verifying that the CD4 and CD8 T-cell responses were MHC class I and II-restricted.

Further investigation utilized blood from uninoculated steers and revealed the same CTL response could be elicited entirely ex vivo by using two rounds of stimulation of cDC and MoDC pulsed with MMP (6). The proliferative and CTL responses were reduced if either CD4 or CD8 T cells were depleted from mdPBMC before DC were pulsed with MMP. This finding suggested the co-presence of CD4 and CD8 T cells with MMP pulsed DC is required to elicit the maximum proliferative and CTL responses. The present study was conducted to explore this possibility in greater detail. As reported, the data provide evidence showing MMP is taken up by DC (cDC and MoDC) via the exogenous route and processed for Ag presentation by MHC class I and MHC class II molecules. Depletion of either CD4 or CD8 T cells in mdPBMC prior to Ag stimulation reduced the proliferative and CTL responses to MMP processed and presented by DC. The proliferative and CTL responses were blocked in the presence of antibodies to either or both MHC I and MHC II. The data provide evidence that the generation of primary CTL responses to MMP require simultaneous cognate recognition of antigenic peptides presented by MHC class I and II molecules. The findings presented in the study reveal a critical component of the immune response to peptide-based vaccines.

## Material and methods

### Animals

Three Holstein steers were obtained from the *Map*-free Washington State University (WSU) dairy herd in 2017. The steers were kept in an open feed lot and used as a source of blood to conduct the ex vivo studies on the immune response to MMP. Steers were maintained by the WSU animal care staff, and all protocols were approved by the WSU Institutional Animal Care and Use Committee (ASAFs 3360 and 04883).

### Preparation of MMP and Map K10

The full length MMP used for antigen presentation by antigen presenting cells (APC) is encoded by *MAP2121c* in the K-10 genome (11). It was expressed in ClearColi as a maltose-binding protein fusion for purification (12). Cultures of *Map* K10 were prepared from single colonies and used to inoculate Middlebrook 7H9 broth flasks (Difco, BD biosciences, USA) supplemented with 6.7% para-JEM GS (Trek Diagnostic Systems, OH), 2 μg/mL mycobactin J (Allied Monitor, MO, USA), and 0.05 % Tween 80 (Sigma-Aldrich Corp.) (9, 13). The cultures were expanded on a shaking stand at 37°C. When the broth cultures reached an OD600 of 0.6 to 0.8, master stocks were prepared in 1.5 mL micro-centrifuge screw-cap tubes for immediate use in each experiment. To ensure a single-cell suspension, bacterial stocks were disaggregated by passages through a 26-gauge needle three times, and bacterial numbers were estimated based on the final OD600 values (13).

### Blood processing for cell separation and culture

The flow diagram in Figure 1 illustrates the protocols used to conduct studies with cells obtained from the three *Map-*negative Holstein steers. As illustrated in Fig. 1, peripheral blood mononuclear cells (PBMC) were prepared by density gradient centrifugation using Ficoll-Hypaque (0.077). One portion of the PBMC was used to generate MoMΦ for use in the viability assay as described below. The second PBMC portion was labelled with magnetic microbeads coated with a cross-reactive anti-human CD14 mAb to isolate monocytes per the manufacturer’s instructions (Miltenyi Biotec) (14). The average purity of isolated CD14^+^ cells was greater than 98%, as determined by FC analysis using an anti-bovine CD14 mAb, CAM36A (15, 16). Monocytes (2 × 10^6^) were added to wells of six well culture plates and cultured in 3 mL of complete culture medium (cRPMI) [RPMI-1640 medium with GlutaMAX^TM^ (Life Technologies, CA) supplemented with 10 % calf bovine serum (CBS), 1 mM β-mercaptoethanol, 100 units/mL of penicillin G, and 100 μg/mL of streptomycin sulfate] in the presence of a DC growth cocktail containing bovine GM-CSF and IL-4 (Kingfisher Biotech, MN). On the third day, 1.4 mL of the medium was replaced with 1.8 mL of fresh medium containing the cocktail. The cultures were incubated for an additional three days to obtain MoDC.

**FIGURE 1.**
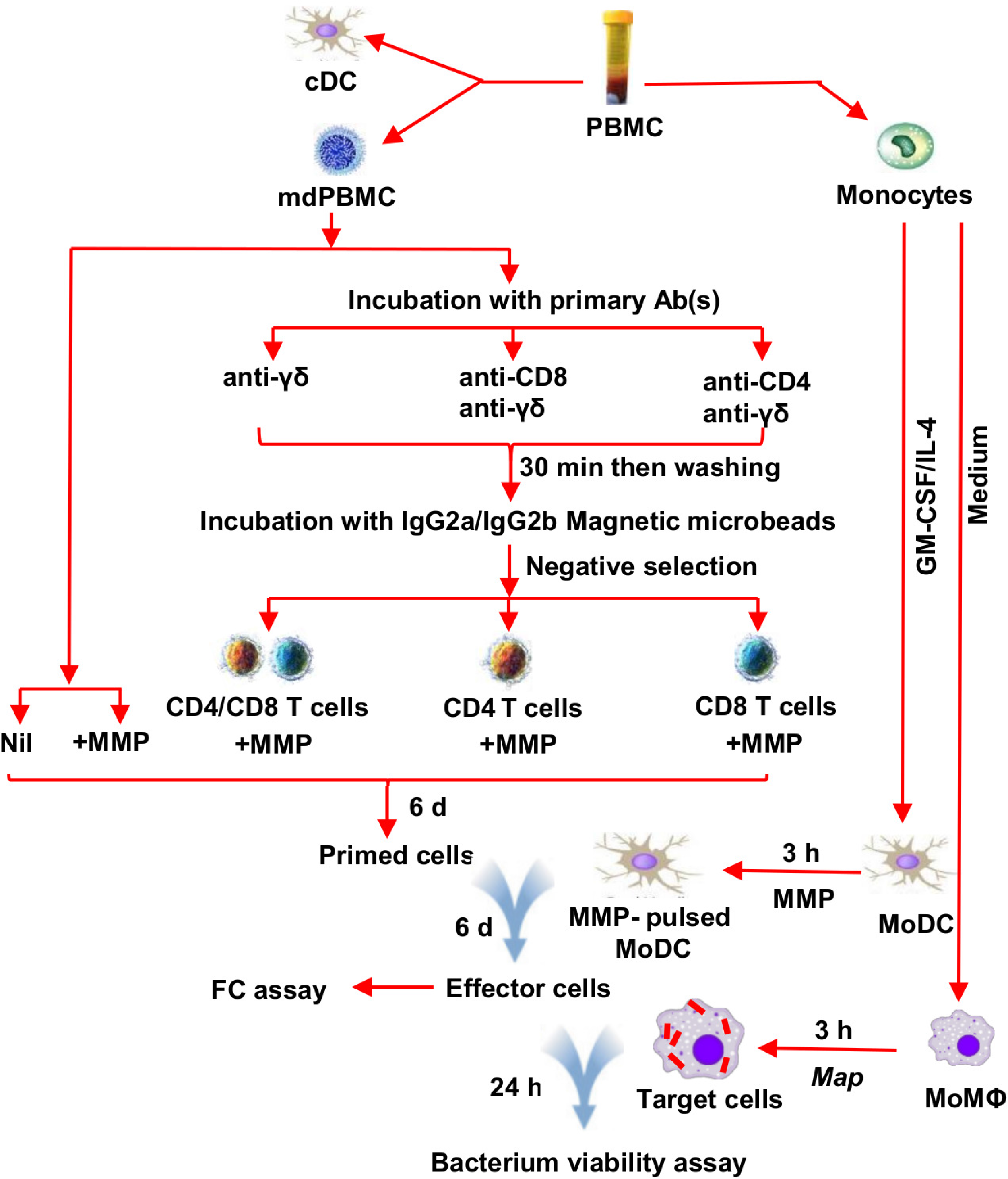
Flow diagram illustrating preparation of mdPBMC for analysis of the immune response to MMP ex vivo. See methods for detail.

The mdPBMC were initially incubated for 30 minutes at 37°C, 5% CO2 with the following combinations of the primary mAbs with no azide (1 ug each/ 10^6^ cells): Anti-CD8 and anti- γδ T cells; Anti-CD4 and anti- γδ T cells; and anti- γδ T cells. After incubation, cells were washed three times with warm RPMI to remove unbound antibodies.

The three sets of primary mAb-treated mdPBMC were then incubated for 15 minutes at 4°C with anti-mouse IgG2a+b microbeads per the manufacturer’s instructions (Miltenyi Biotec). After incubation, the cells were washed two times with cold MACS buffer (a solution containing phosphate-buffered saline (PBS), pH 7.2, 0.5% bovine serum albumin (BSA), and 2 mM EDTA) to remove the unbound microbeads, and then re-suspended in the same buffer. For magnetic separation, LS columns were placed in the magnetic field of a MACS Separator and rinsed with 3 mL of MACS buffer, and cell suspensions were loaded onto the wet columns. Flow-through containing the following three unlabeled cell fractions of mdPBMC were collected (negative selection): CD4 T cells, CD8 T cells and CD4/CD8 T cells. Unseparated mdPBMC were also maintained for use as positive and negative control wells.

The three mdPBMC fractions and unseparated mdPBMC were subjected to two rounds of antigenic stimulation using MMP. To conduct the first round of stimulation, cells were distributed in the 6-well culture plate in duplicate (2 × 10^6^/mL in 5 mL of cRPMI). MMP (5 μg/mL) was added to each well and incubated for 6 days at 37°C, 5% CO2 to allow MMP processing and presentation by cDC present in the mdPBMC. To conduct the second round of stimulation after 6 days of culture, MMP (5 μg/mL) was added to the cultures of MoDC and incubated for 3 hours at 37°C, 5% CO2. The adherent MoDC were then carefully washed 3 times with warm RPMI to remove the cell-free MMP. The primed cells were collected, washed twice with warm RPMI, and added to their respective autologous MoDC pulsed with MMP (2 × 10^6^/mL in 5 mL of cRPMI). After six additional days of culture, the cells were collected and used in FC and the *Map* viability assay as described below. A portion of the unseparated mdPBMC was maintained as a negative control with no antigen stimulation during the two weeks of cell culture.

### Viability assay

Control and antigen-stimulated mdPBMC and CD4 T-cell, CD8 T-cell, and CD4/CD8 T-cell fractions were used as effector CTLs in the viability assay.

### Generation of MoMΦ for use as target cells

As mentioned above, one portion of fresh PBMC was re-suspended in cRPMI transferred into 150 mm tissue culture plates and incubated overnight. The majority of the non-adherent cells were then removed the following day. The adherent cells were cultured in fresh medium for six days then brought into suspension by incubation on ice in the presence of EDTA in PBS (4 mL EDTA [250 mM stock in H2O], 5 mL CBS, 91 mL PBS). The MoMΦ were distributed into six well culture plates (2 × 10^6^ cells/ well) and cultured for an additional six days, and then used as target cells in the viability assay.

### Infection of target cells with Map K10

MoMΦ were infected with *Map* K10 at a multiplicity of infection (MOI) of 10:1 (2 × 10^7^ *Map* to ~2 × 10^6^ MoMΦ/well) in antibiotic-free cRPMI. Culture plates were centrifuged at 700 × *g* for five minutes, then incubated at 37°C, 5% CO_2_ for 3 hours. Extracellular bacteria were removed by washing five times with warm, antibiotic-free RPMI using gentle suction to avoid detaching adherent MoMΦ. Two wells from each of the respective sets of 6 wells, containing *Map* infected MoMΦ, were used as controls, without addition of primed or unprimed preparations of mdPBMC.

### Incubation of effector T cells with infected target cells

Stimulated and control mdPBMC and T-cell fractions were collected and added to the preparations of infected MoMΦ. Co-cultures were incubated for 24 hours at 37°C, 5% CO_2_. Non-adherent cells were collected, then adherent cells were detached and collected as described above. Finally, collected adherent and non-adherent cells were recombined for analysis of *Map* viability.

### Cell lysis

Following collection, cells were lysed by adding 2 mL of 0.01% saponin in H_2_O and incubated at 37°C for 15 minutes. The cell lysates were centrifuged for 30 minutes at 4,500 rpm to pellet the bacteria. The supernatants were discarded and the pellets re-suspended in 1 mL H_2_O and transferred into micro-centrifuge tubes, then centrifuged at 14,000 rpm for 10 minutes. The supernatants were discarded, and the pellets re-suspended in 400 μl of H_2_O in 1.5 mL translucent Eppendorf tubes and stored at −20°C until used.

A set of controls was prepared from known mixtures of live and dead *Map* K10. This set of controls covered the dynamic range for detection of live vs dead *Map* obtained from infected MoMΦ before and after incubation with CTL. Aliquots of *Map* mixed in five ratios, 100% live, 75% live/25% killed, 50% live/50% killed, 25% live/75% killed, and 100% killed, were prepared to obtain 2 × 10^7^ total *Map* in each aliquot, added to the cultures of MoMΦ at a MOI of 10, and incubated for 3 hours as described previously (6). The cultures were then washed to remove free bacteria. Adherent cells were collected and transferred into new 15 mL tubes. 10^7^ fresh mdPBMC were added for each tube and all the cells were mixed followed by centrifugation. The cell pellet in each tube was lysed with saponin as described above, and lysates stored at −20°C until use.

### PMA treatment, DNA extraction and qPCR

Propidium monoazide (PMA) treatment of the cell lysates was carried out as previously described (6). Briefly, 1 μl of 20 mM PMA working stock solution was added to 400 μl of each previously prepared cell lysates to reach a final dye concentration of 50 μM. The translucent PMA-treated tubes were incubated at room temperature for five minutes in the dark on a rocker. The tubes were then placed in a plastic tray prepared with a frozen ice pack wrapped in aluminum foil. The tray was then placed on a rocking platform to ensure continuous mixing during light exposure. Light exposure was performed for five minutes using a halogen lamp with a 650 W bulb placed at a distance of ~ 20 cm from the tubes. Cells were subsequently harvested by centrifugation at 10,000 × *g* for five minutes. Supernatants were discarded, and the cell pellets processed for DNA isolation (17).

DNA extraction was performed according to the protocol for Gram-positive bacteria using DNeasy^®^ Blood and Tissue kit (Qiagen, USA) following enzymatic lysis to facilitate breakdown of the *Map* cell wall as described by Park et al. (17). TaqMan Quantitative Real-Time PCR, targeting the single copy F57 gene specific for *Map* (F57 qRT-PCR) was used to determine the activity of intracellular *Map* killing as described by Kralik et al. (18) and Abdellrazeq et al. (6). The reaction was performed according to Schönenbrücher et al. (19) using a StepOnePlus Real-Time PCR machine (Applied Biosystems, CA). *Map* gDNA prepared from pure culture was used to generate a standard curve with the F57 probe, made with 8 dilutions starting with 4 × 10^7^ copies down to 4 copies. The sequences of the primers and probes were the same as previously described (17). The total reaction volume was 25 μL including 5 μL of the DNA sample, and reactions were run for 40 cycles. The mean values of the cycle threshold (C_T_) were analyzed using StepOne Software v2.1 (Applied Biosystems, CA).

### MHC blocking

Three identical sets of unseparated mdPBMC cultures were prepared in the presence of mAbs specific for either or both MHC I and MHC II (0.5 μg/ml) (Table 1). Cultures were subjected to two rounds of stimulation with MMP as described above, and cell activation and proliferation assessed using FC. The resultant cultures of cells were then incubated with infected MoMΦ for 24 hours. The cells were collected and processed to determine the CTL activity in each preparation of cells using the bacterium viability assay as described.

**Table I.**
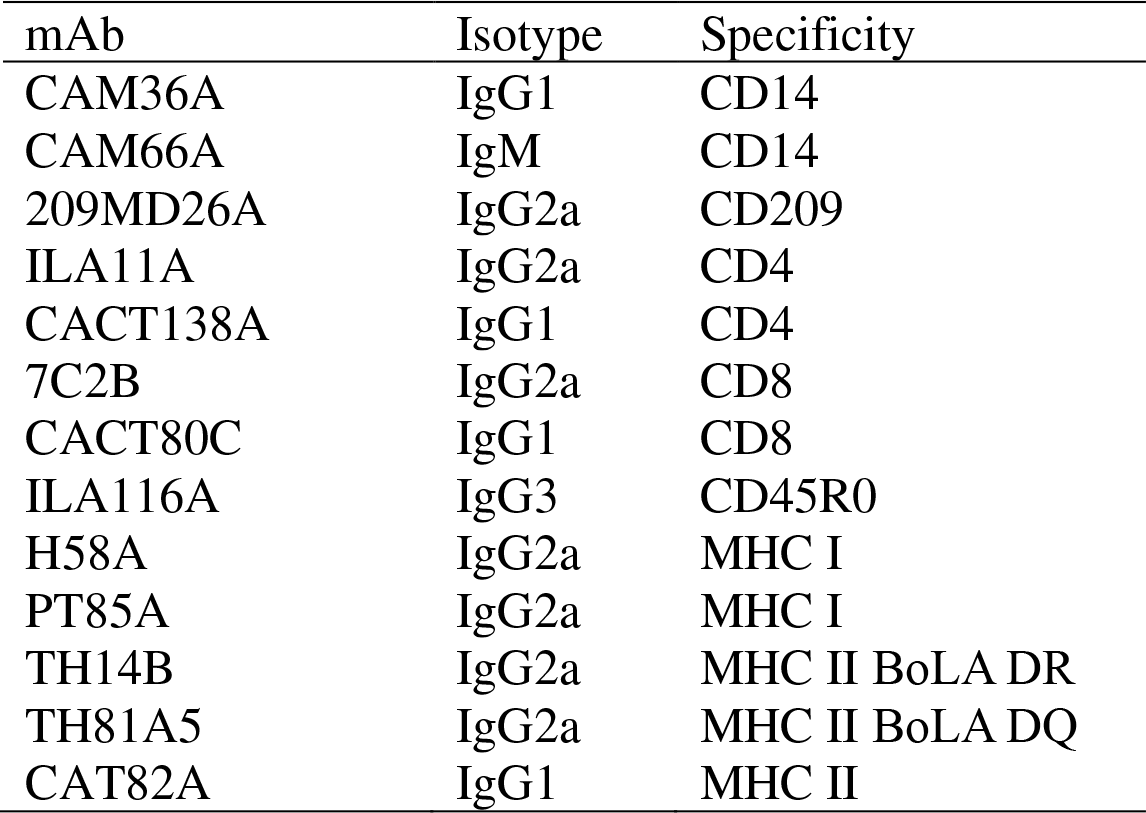
Monoclonal antibodies used in the present study.

| mAb | Isotype | Specificity |
| --- | --- | --- |
| CAM36A | IgG1 | CD14 |
| CAM66A | IgM | CD14 |
| 209MD26A | IgG2a | CD209 |
| ILA11A | IgG2a | CD4 |
| CACT138A | IgG1 | CD4 |
| 7C2B | IgG2a | CD8 |
| CACT80C | IgG1 | CD8 |
| ILA116A | IgG3 | CD45R0 |
| H58A | IgG2a | MHC I |
| PT85A | IgG2a | MHC I |
| TH14B | IgG2a | MHC II BoLA DR |
| TH81A5 | IgG2a | MHC II BoLA DQ |
| CAT82A | IgG1 | MHC II |

### Flow cytometric analysis

After the second round of antigen stimulation, cells were washed once in PBS/ACD, centrifuged, re-suspended in serum-free RPMI and counted. For antibody labeling, cells were distributed into 96-well polystyrene V-bottom microplates (10^6^ cells/well). Combinations of mAbs (Table 1) obtained from the WSU Monoclonal Antibody Center (WSUMAC) were used for labeling as previously described (20). Data were collected on a modified FACS Calibur DxP8 Analyzer equipped with a FlowJo operating system (Cytek Biosciences Inc. Fremont, CA) and analyzed with FCS Express software (DeNovo Software, Glendale, CA) (15). The gating strategy used to collect the data is shown in Fig. 2. Side scatter (SSC) and forward scatter (FSC) were used to identify small and large lymphocytes. FSC-Height vs FSC-Width (FSC-H vs FSC-W) was used to exclude doublets. Selective electronic gating was used to isolate CD4 and CD8 T cells for determination of their activation status.

**FIGURE 2.**
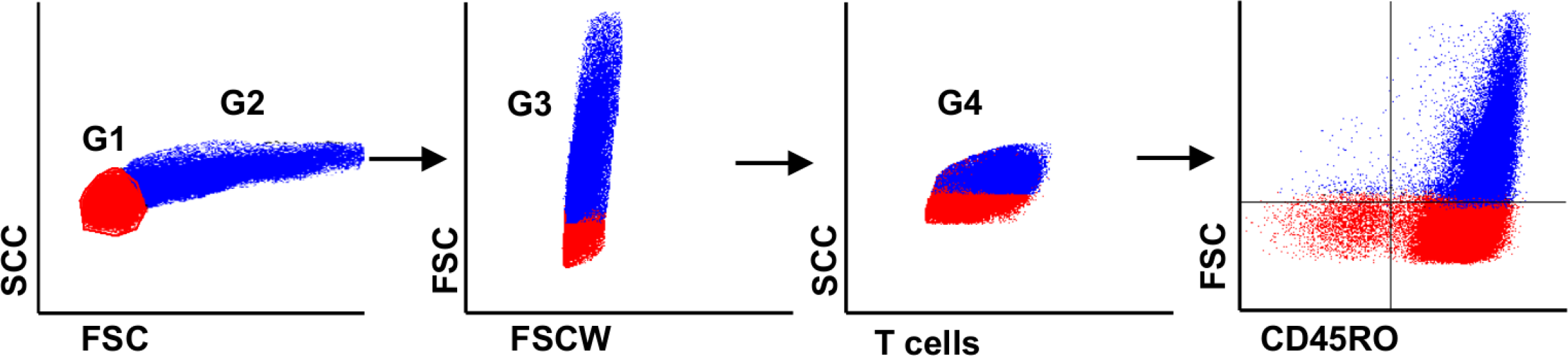
Flow cytometric gating strategy. Gates were used to isolate CD4 and CD8 T cell subsets for analysis of their proliferative response following stimulation with cDC and MoDC pulsed with MMP. **(A)** Two light scatter parameters, side scatter (SSC) vs forward scatter (FSC) were used to identify the small lymphocytes (G1, color coded red) and large activated lymphocytes (G2, color coded blue) based on size and granularity. (B) A pulse geometry gate (FSC-H vs FSC-W) was placed on single cells (G3) to exclude doublets. (C) A fourth gate (G4) was used to isolate CD4 and CD8 T cells to determine their activation status. For data analysis, FSC vs CD45R0 (a memory T cell marker) was used to distinguish non-activated memory cells (red) from activated memory cells (blue) proliferating in response to stimulation with MMP. (D) Only the activated memory cells (upper right quadrant in the dot plot) were considered for statistical analysis.

### Statistical methods

Data were imported into SAS software (SAS for Windows, version 9.3) for statistical analysis and graphical presentation. The data were analyzed using a mixed modeling procedure (PROC GLIMMIX). For proportional response data (proportion activated T cells), statistical models included the main fixed effects of MMP stimulation (MMP, diluent control) and T cell type (CD4, CD8), the interaction term of these effects, and was based on the binomial response distribution and Kenward-Roger degrees of freedom approximation. These experiments were considered to be of heirarchical design; the corresponding statistical models thus included random residuals defined by T cell types nested within each subject (blood donor steer), and an unstructured (Cholesky) covariance matrix. Multiple comparisons were adjusted using the method of Holm-Tukey (overall α = 0.05).

For C_T_ response data (qPCR estimation of intracellular *Map* killing), statistical models included the single main fixed effect of manipulating the mdPBMC context of MMP stimulation (manipulation of T cell context or manipulation of MHC I and II context, each with controls). These analyses were based on the Poisson response distribution and Kenward-Roger degrees of freedom approximation, and included a random effect of residuals defined by subjects and using the default variance component structure of the covariance matrix. Under these experimental scenarios, only comparisons of interest were made utilizing the Dunnett method of adjustment for multiple comparisons (overall α = 0.05).

## Results

### The proliferative response of mdPBMC is reduced if either CD4 or CD8 T cells are depleted before stimulation with MMP

The experiments were conducted to verify and gain further insight into the effect of CD4 and CD8 depletion on CTL development following MMP stimulation. Negative selection was used to deplete either CD4 or CD8 T cells from mdPBMC. The depletion strategy included use of a mAb to the δ chain of the γδ TCR to exclude any potential effect of γδ T cells on the proliferative response to MMP. Previous studies established that NK cells do not contribute to the proliferative response to MMP (6). Little difference could be detected between the untreated positive control preparation of mdPBMC and preparations depleted of either CD4 or CD8 T cells after one round of stimulation with MMP (not shown). However, clear differences were evident after two rounds of stimulation (Fig. 3).

**FIGURE 3.**
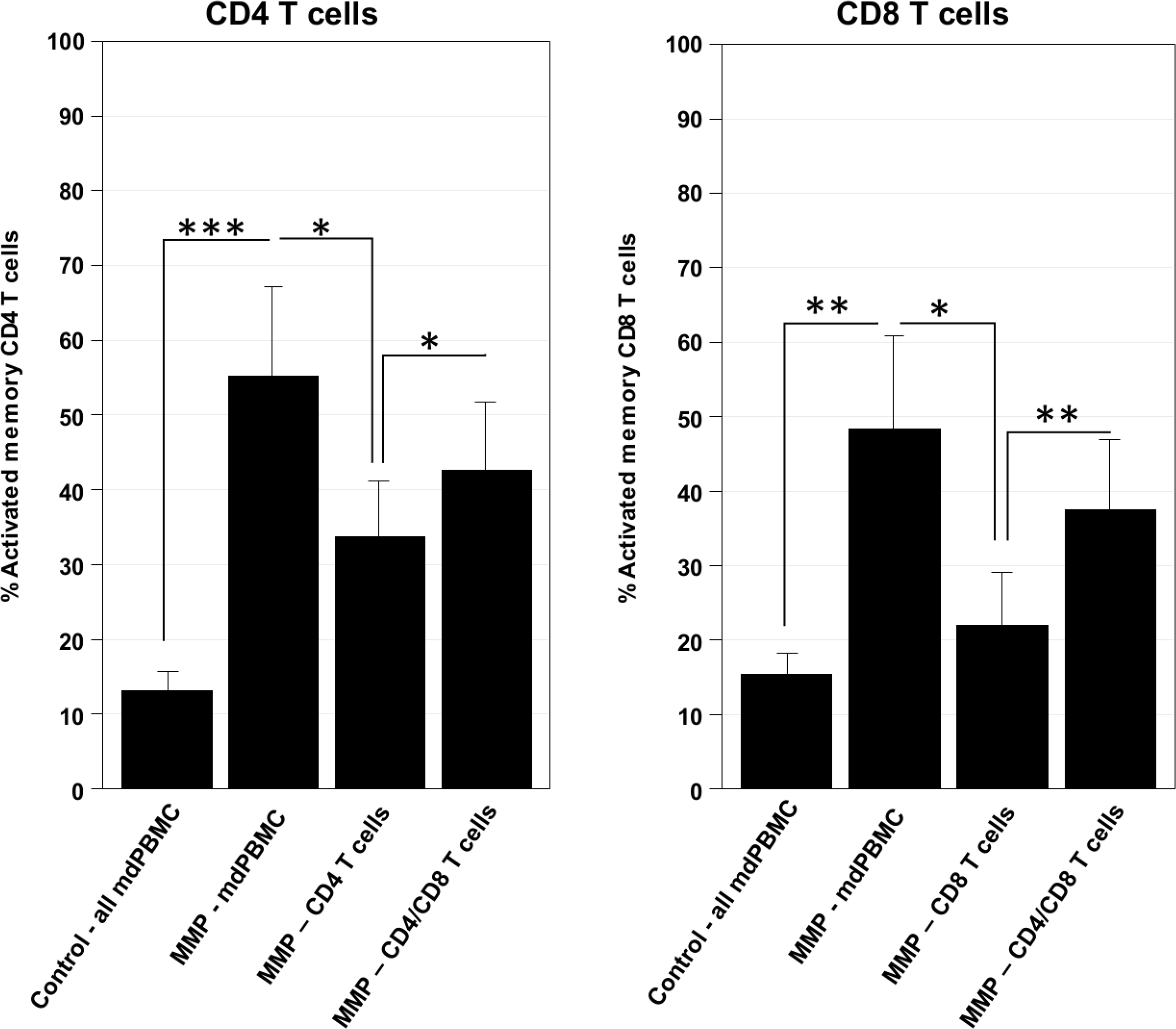
The role of cellular context on CD4 and CD8 T cell activation by MMP. Two-rounds of stimulation of unseparated mdPBMC (all mdPBMC) with MMP increased the percentages of activated CD4 T cells and CD8 T cells (control vs MMP). The percentages of activated CD4 T cells and CD8 T cells after stimulation were not significantly reduced if the mdPBMC were first depleted of γδ T cells (cellular context: CD4/CD8 T cells versus all mdPBMC). The percentage of activated CD4 T cells after stimulation was reduced if the mdPBMC were depleted of γδ T cells and CD8 T cells (CD4 T cells versus all mdPBMC, and versus CD4/CD8 T cells). The percentage of activated CD8 T cells after stimulation was similarly reduced if the mdPBMC were first depleted of γδ T cells and CD4 T cells (CD8 T cells versus all mdPBMC, and versus CD4/CD8 T cells). Data shown are the least squares means and standard deviations for experiments on blood collected from 3 steers. Significance symbols represent *P*-values adjusted for all pairwise comparisons such that: *, *P*_*adj*_ < 0.05; **, *P*_*adj*_ < 0.01; ***, *P*_*adj*_ < 0.001.

mdPBMC incubated with MMP had significantly larger (*F*_MMP_=168.06, *P*=0.0002) proportions of activated CD4 T cells (*P_adj_*=0.0005) and activated CD8^+^ T cells (*P_adj_*=0.0013) than mdPBMC incubated without MMP. Significant differences were not detected between the proportions of activated CD4 and CD8 T cells within the mdPBMC (*F*_CD_=0.7277, *P*=0.7277) nor in the change induced by MMP (*F*_MMP*CD_=2.61, *P*=0.1816).

The cellular context during incubation with MMP did have a significant effect on the proportions of activated T cells (*F*_Context_=44.68, *P*=0.0053). Post-hoc comparisons were limited to within each cell type since the effects on the two phenotypes of T cells were parallel (see Fig. 3) and only marginally different in magnitude (*F*_Context*CD_=13.23, *P*=0.0308). The proportion of activated CD4 T cells in mdPBMC depleted of both CD8 T cells and γδ T cells was significantly less than the proportions present in whole mdPBMC (*P_adj_*=0.0327) and mdPBMC depleted of only γδ T cells (*P_adj_*=0.0225). The proportion of activated CD4 T cells in mdPBMC depleted of γδ T cells was not significantly different than that in whole mdPBMC (*P_adj_*=0.0542).

Similarly, the proportion of activated CD8 T cells in mdPBMC depleted of both CD4 T cells and γδ T cells was significantly less than the proportions within whole mdPBMC (*P_adj_*=0.0131) and mdPBMC depleted of only γδ T cells (*P_adj_*=0.0022). The proportion of activated CD8 T cells in mdPBMC depleted of only γδ T-cells was not significantly different than that in whole mdPBMC (*P_adj_*=0.0786).

### The CTL activity of mdPBMC is reduced if either CD4 or CD8 T cells are depleted before stimulation with MMP

In conjunction with experiments to investigate the effects of T-cell depletion on the proliferative response to MMP, one set of cell preparations was used to determine the effects of depletion on CTL activity against intracellular *Map* in target MoMΦ (Fig. 4). Manipulation of cellular context did significantly affect killing of intracellular bacteria (as estimated by C_T_; *F*=16.40, *P*<0.0001). Fig. 4 depicts the outcomes of this experiment relative to a standardized scale of intracellular killing whereas statistical comparisons to the maximal killing produced by whole mdPBMC 24-hours post stimulation with MMP are shown in Fig. 5. The intracellular killing of *Map* in MoMΦ co-cultured with MMP-stimulated whole mdPBMC was significantly greater than the control conditions (vs. MoMΦ at 0-hours and at 24-hours post-stimulation, each *P_adj_*<0.0001; vs. MoMΦ-control at 24-hours post-stimulation, *P_adj_*=0.0005). A significant reduction in intracellular killing of *Map* was not detected when only γδ T-cells were depleted from mdPBMC prior to MMP stimulation (*P_adj_*=0.4682), but was detected with the additional depletion of CD4 T cells (*P_adj_*=0.0457) or CD8 T cells (*P_adj_*=0.0089).

**FIGURE 4.**
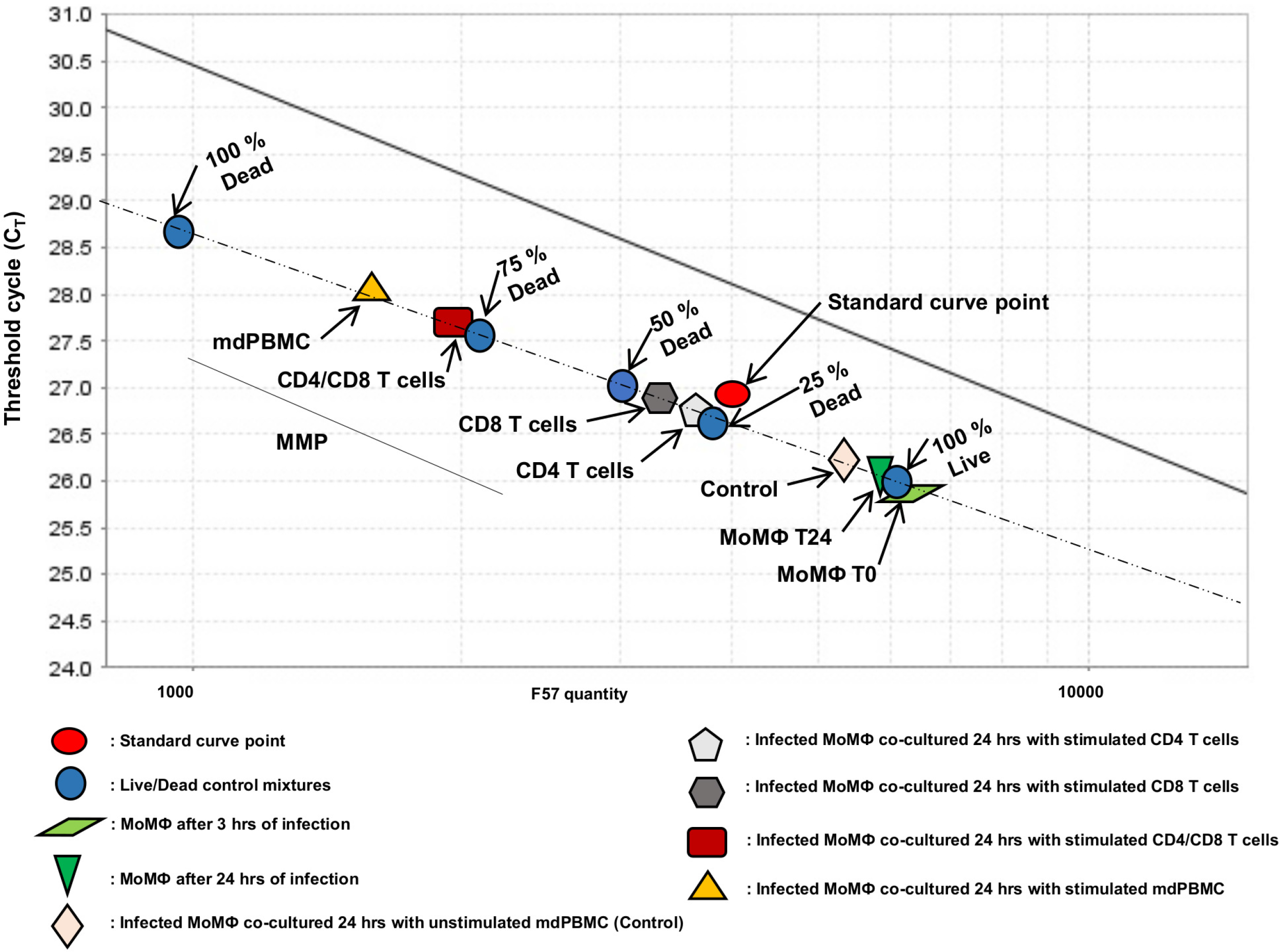
The role of cellular context during stimulation by MMP on intracellular killing of *Map*. Illustration of data presented on a representative standard DNA curve. The number of live bacteria were estimated by qPCR (C_T_ values). Standards of live:dead *Map* bacteria are labeled above the data (blue connectors). Note that the relationship of C_T_ values with live bacterial content (log of F57 quantity) is inverse and linear within the effect range of interest. Experiment test conditions (MMP stimulation and cellular context of mdPBMC) are noted below the data (independently colored shapes).

**FIGURE 5.**
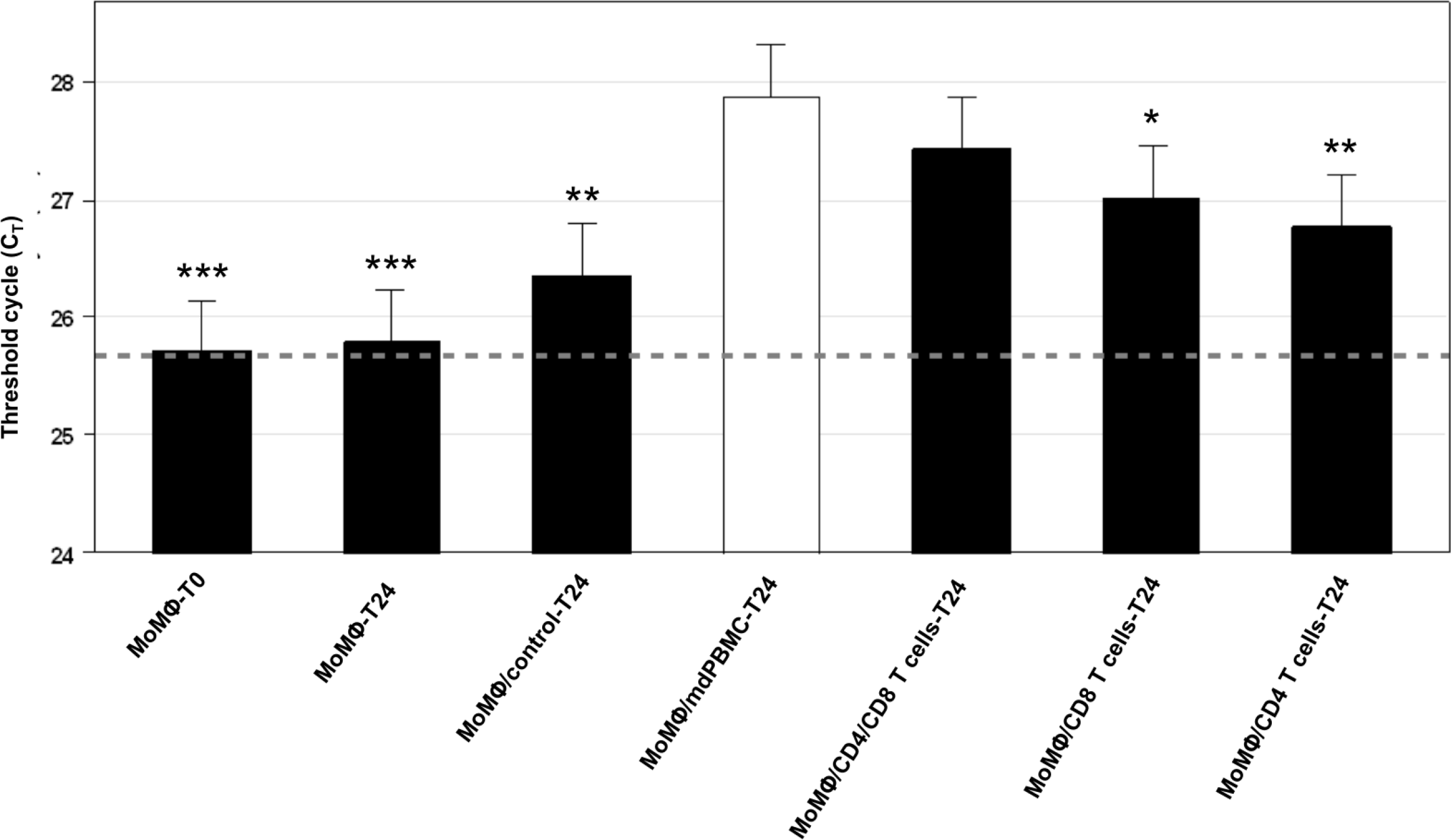
Cumulative C_T_ values of intracellular killing of *Map*. In comparison to control conditions (left most solid bars), significant killing of *Map* (reduced live bacteria = higher value) was detected after infected MoMΦ were co-cultured 24-hours with unseparated mdPBMC stimulated by MMP (open bar). Manipulation of cellular context during stimulation of mdPBMC with MMP (rightmost solid bars) did not significantly reduce intracellular killing with depletion of only gamma-delta T cells (cellular context: “CD4/CD8 T cells-T24”) but was significantly reduced if mdPBMC were also depleted of CD4 T cells (CD8 T cells-T24) or CD8 T cells (CD4 T cells-T24). Data shown are the least squares means and standard deviations for experiments on blood collected from 3 steers. Significance symbols represent *P*-values adjusted for multiple comparisons to the condition of maximum killing activity (open bar) such that: *, *P*_*adj*_ < 0.05; **, *P*_*adj*_ < 0.01; ***, *P*_*adj*_ < 0.001.

### Development of CD8 CTL to Map is inhibited by mAb blockade of MHC class I and/or class II molecules

Depletion experiments confirmed the requirement of both CD4 and CD8 T cells during priming by DC for the generation of significant anti-*Map* CTL activity ex vivo. However, depletion experiments did not reveal whether development of CTL activity required cognate recognition of MMP epitopes presented on DC MHC I and MHC II molecules by CD4 and CD8 T cells during antigenic stimulation. Our previous studies of the recall response to *Map/*Δ*relA* and MMP revealed the recall response was blocked in the presence of mAbs to both MHC I and MHC II molecules (10), but the effect of individual MHC class I or class II molecule blockade was not investigated. To complete the last set of experiments in this study, mdPBMC were stimulated with MMP in the presence of mAbs specific for either or both MHC I and MHC II molecules (Table 1). To maintain consistency with the initial studies, two mAbs were used for MHC I blockade and two mAbs (one specific for the bovine orthologue of HLA-DR, one specific for the bovine orthologue of HLA-DQ) were used for MHC II blockade. The effect of MHC molecule blockade on cell proliferation and the development of CTL activity against intracellular *Map* was significant (*F*=29.05, *P*<0.0001). The intracellular killing of *Map* by CTL in whole mdPBMC was significantly greater (all *P_adj_*<0.0001) than the negative control conditions and to that observed in cultures containing mAbs to either MHC I or MHC II molecules as well as in the presence of both MHC I and MHC II mAbs (Fig. 6).

**FIGURE 6.**
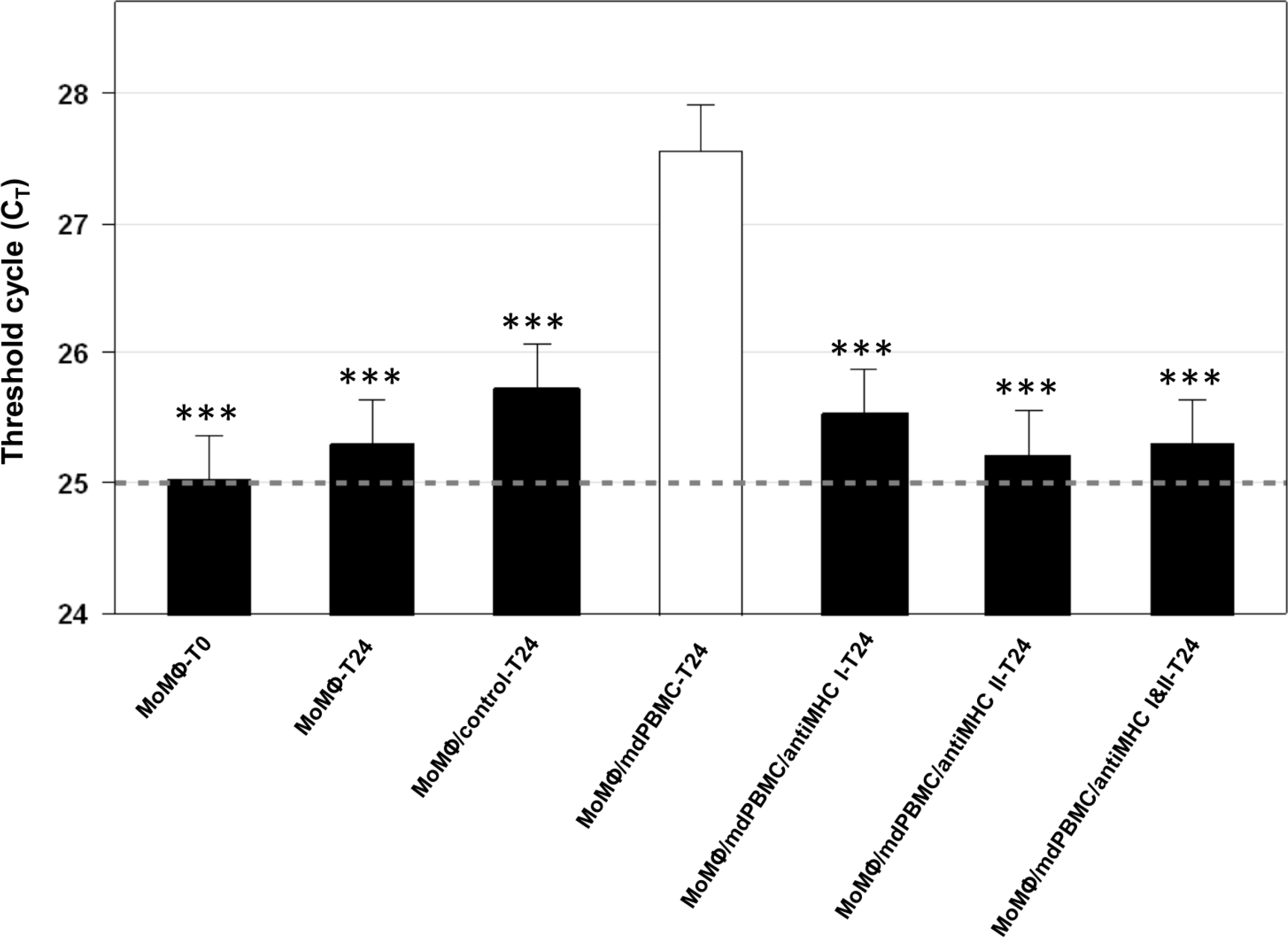
The effect of anti-MHC antibodies present in mdPBMC during stimulation by MMP on intracellular killing of *Map*. In comparison to control conditions (leftmost solid bars), significant killing of *Map* (reduced live bacteria = higher C_T_ value) was detected after infected MoMΦ were co-cultured 24-hours with unseparated mdPBMC stimulated by MMP (open bar). The presence of antibodies to MHC I, MHC II, or both MHC I and MHC II during stimulation of mdPBMC with MMP (rightmost solid bars) significantly reduced cell proliferation and intracellular killing of *Map*. Data shown are the least squares means and standard deviations for experiments on blood collected from 3 steers. Significance symbols represent *P*-values adjusted for multiple comparisons to the condition of maximum killing activity (open bar) such that: *, *P*_*adj*_ < 0.05; **, *P*_*adj*_ < 0.01; ***, *P*_*adj*_ < 0.001.

## Discussion

Extensive studies have been conducted to elucidate the mechanisms regulating development of CD8 CTL activity against viral and bacterial pathogens, intracellular parasites, and cancers with the long-term objective of vaccine development. Cumulative studies have shown DC play a central role in stimulating CTL activity through cross presentation of antigenic epitopes presented in context of MHC I molecules (reviewed in (21, 22)). What has not been fully explained in these studies is the role of CD4 T cells in the development of CD8 CTL activity. Indeed, reports on the role of CD4 T cell help seems to vary greatly between different model systems. Primary CTL response development to highly inflammatory targets, such as whole organism or bacterial membrane-based vaccines, are reported to be CD4 T cell independent (23, 24), while CD4 T cell help during priming are reported to be necessary for the development of a functional memory CTL response to these antigens (25–28). CD4 T-cell help has also been reported to be necessary for activation of the CTL recall response. In contrast, the generation of both primary and recall CD8 CTL responses to poorly antigenic targets, including peptide immunogens and neoplastic cells, have been reported to require CD4 T cell help during priming and recall responses (Ridge et al., 1998, Bennett et al., 1998).

The mechanism of action of CD4 T cell help in these systems has not been clearly elucidated. Some studies have suggested CD4 T cells may play an indirect role and that interaction of CD154 (expressed on CD8 T cells) with CD40 (expressed on APC) might be the triggering event that initiates the primary activation and secondary expansion of CD8 T cells (reviewed in (29–31)). Results from other studies have suggested that cognate recognition of antigens presented in the context of DC MHC class II and class I molecules, coupled with subsequent stimulation by DC through ancillary receptor-ligand interactions, are key steps in priming CTL responses (32, 33). Although complex inbred mouse models have provided insight into the events associated with the generation of CTL in vivo, additional information is still needed to fully detail the events regulating development of CTL responses in outbred species, like cattle and humans. The present studies were conducted using an outbred bovine model system to characterize the CTL response to a candidate *Map* peptide-based vaccine.

Key to the development of our model system was the finding that CD209 is uniquely expressed on cDC in blood and MoDC in cattle (10). Due to the size of cattle, access to large volumes of blood obviated the difficulty in obtaining sufficient quantities of cDC for comparative studies with MoDC (10). The development of a bovine DC-lymphocyte culture system enabled dissection of primary and recall CTL immune responses to antigenic peptides presented in context of MHC I and MHC II molecules under defined conditions. The *Map*/Δ*relA* major membrane protein antigen, MMP, provided a model peptide antigen to study factors affecting development of CTL to a candidate peptide vaccine. Finally, development of a bacterium viability assay provided a way to study the mechanisms used for the intracellular killing of *Map* in infected target cells (6).

Using a *Map/ΔrelA-*vaccinated steer, we first used our model system to dissect the ex vivo recall response to *Map/ΔrelA* and MMP. The studies demonstrated that the main cell subsets proliferating in response to stimulation with *Map/ΔrelA* or MMP-pulsed DC were CD4 and CD8 T cells. The responses were MHC-restricted. Killing activity was mediated primarily by CD8 T cells through the perforin-granzyme B pathway (6). A preliminary CD4 T cell depletion study indicated development of CTL activity was reduced if CD4 T cells were removed from cultures before stimulation with Ag-pulsed DC (6). The data provided support for the supposition that CD4 T cell help is essential for initiation of a functional CD8 T cell recall response to *Map/ΔrelA* and MMP.

Since CD4 T cell help during priming is often observed for the generation of a primary CTL response to non-inflammatory targets in mice, we conducted the present ex vivo study to determine if this observation was the same in regard to the *Map/ΔrelA* and MMP candidate vaccines. In this study, we found the primary CD8 CTL responses were significantly diminished in cultures depleted of CD4 T cells prior to antigenic stimulation by DC. We also found that CD8 CTL responses were blocked in whole mdPBMC cultures in the presence of mAbs specific for MHC class I and MHC class II molecules. Importantly, we observed there was complete blockade of CD4 and CD8 T cell activation in the presence of mAbs to either MHC I or MHC II alone. The data presented here provide evidence that cognate CD8 and CD4 T cell recognition of antigenic peptides presented on MHC I and MHC II molecules is essential for development of a functional primary CTL response to MMP.

The importance of CD4 and CD8 T cell cognate antigen recognition to the development of a functional primary CTL response to a bovine pathogen was previously demonstrated in one other study using an ex vivo *Theileria parva* culture system (34). In this study, *T. parva*-infected lymphocytes, which express MHC class I and class II molecules, were used as APC, and were cultured in the presence of *T. parva-*naïve CD8 T cells, with or without primed CD4 T cells. *T. parva-*specific CD8 CTL developed only in cultures that contained primed CD4 T cells. Furthermore, if the primed CD4 T cells were cultured in the same plate, but separated from the APC and CD8 T cells by a semi-permeable membrane, no primary CD8 CTL response to *T. parva* developed, indicating that direct cell-cell contact between APC, CD4 T cells, and CD8 T cells is required for the development of a primary CTL response in this system. This study, like the present study, provides evidence of the importance of simultaneous cognate antigen recognition by CD4 and CD8 T cells for the development of a functional CTL response in the cattle.

In summary, analysis of the immune response to a *Map/relA* deletion mutant and a candidate peptide-based vaccine for *Map* in cattle have provided insight into factors regulating the development of CTL to an intracellular pathogen that are of universal importance. Deletion of *relA* disrupted the pathways used by *Map* to dysregulate the immune response and allowed for the development of an immune response that cleared the infection with the mutant. Analysis of the recall response revealed vaccination led to development of a CD8 CTL response that targeted a membrane protein, MMP. Analysis of the entire immune response to MMP ex vivo revealed simultaneous cognate recognition of antigenic peptides by CD4 and CD8 T cells, presented by DC pulsed with MMP is essential for generation of CTL against *Map*. Blocking of Ag presentation by mAbs to either MHC I or MHC II molecules blocked the proliferative and CTL responses to *Map*. The findings may have revealed an elusive component of the CTL response to pathogens. The ex vivo platform developed to conduct the studies provide an opportunity for further in depth studies in large outbred animal species like cattle and also humans.

## Acknowledgments

The authors wish to acknowledge the excellent technical support and animal care provided by Emma Karol and her staff. The results of this study were presented orally in 2018 Berkeley conference (Mycobacterial Implications in Crohn’s and Chronic Inflammatory Diseases), Lawrence Hall of Science | Berkeley, CA, USA.

## Disclosures

The authors declare that they have no competing interests.

## Funding

Support was provided by the Wash. State Univ. Monoclonal Antibody Center.

## Authors’ contributions

GSA and WCD conceived the study. GSA and WCD participated in the design of the protocol to conduct the studies. JPB participated in the development and use of the *Map* major membrane protein (MMP). GSA conducted the experiments. MME, AHM and VH participated in the conduct of the experiments. GSA and DAS participated in statistical analysis of the data. GSA, LMF, MME, AHM, KTP, WCD, JPB, DAS, and WMC participated in the writing and interpretation of the results. WCD and JPB obtained the funding for the project. WCD oversaw and participated in all aspects of the study. All authors read and approved the final manuscript.

Address correspondence and reprint requests to Dr. William C. Davis, Department Veterinary Microbiology/Pathology, Washington State University, Pullman, WA 99164-7040. E-mail address

## Abbreviations used in this article

mdPBMC: monocyte-depleted peripheral blood mononuclear cells
cDC: conventional dendritic cells
MoDC: monocyte-derived DC
MoMΦ: monocyte-derived macrophages
*Map*: *Mycobacterium avium* subsp. *paratuberculosis*
WT: wild type
MMP: 35 kDa membrane peptide
PMA: Propidium monoazide
qPCR: quantitative PCR
C_T_: cycle threshold.

